# Comparison of Two Individual Identification Algorithms for Snow Leopards after Automated Detection

**DOI:** 10.1101/2022.01.20.477059

**Authors:** Drew Blount, Eve Bohnett, Jason Holmberg, Jason Parham, Sorosh Poya Faryabi, Örjan Johansson, Li An, Bilal Ahmad, Wajid Khan, Stephane Ostrowski

## Abstract

1. Photo-identification of individual snow leopards (*Panthera uncia*) is the primary technique for density estimation for the species. A high volume of images from multiple projects, combined with pre-existing historical catalogs, has made identifying snow leopard individuals within the images cost- and time-intensive. 2. To speed the classification among a high volume of photographs, we trained and evaluated image classification methods for PIE v2 (a triplet loss network), and we compared PIE’s accuracy to the HotSpotter algorithm (a SIFT based algorithm). Analyzed data were collected from a curated catalog of free-ranging snow leopards photographed across years (2012-2019) in Afghanistan and from samples in captivity provided by zoos from Finland, Sweden, Germany, and the United States. 3. Results show that PIE outperforms HotSpotter. We also found weaknesses in the initial PIE model, like a minor amount of background matching, which was addressed, although likely not fully resolved, by applying background subtraction (BGS) and left-right mirroring (LR) methods. The PIE BGS LR model combined with Hotspotter showed a Rank-1: 85%, Rank-5: 95%, Rank-20: 99%. 4. Overall, our results recommend implementing PIE v2 simultaneously with HotSpotter on Whiskerbook.org.

## Introduction

The snow leopard (*Panthera uncia*) is categorized by the International Union for Conservation of Nature (IUCN) as Vulnerable (McCarthy et al., 2017). Population estimates in sampled areas primarily rely on the use of camera-trap technology of individuals identified by their unique spotty phenotypes, in concert with capture-recapture modeling (Jackson et al., 2006; Royle & Young, 2008; McCarthy & Mallon 2016). Abundance estimates for snow leopards have shown to be fraught with errors from camera trap photo misclassification arising from a variety of reasons (Ellis 2018, Johansson et al. 2020), including the manual processing of photo sets, that have become increasingly large with the advent of affordable digital photography (Beery et al., 2019; Falzon et al. 2019; Miguel et al. 2016). Current solutions for reducing the risk of misidentifying images of snow leopards are often resource-intensive, for example, using repetition and multiple observers to manually process large photo sets to limit the risk of false-negative classification (Borchers & Fewster 2016; Choo et al. 2020; Foster & Harmsen 2012; Johansson et al. 2020).

To reduce misclassification errors, as well as time and labor in processing camera trap data, scientists are increasingly turning towards artificial intelligence and computer vision to identify animals through automated image classification by species (Beery et al., 2019; Falzon et al., 2019; Nguyen et al., 2017; Norouzzadeh et al., 2019; Parham et al., 2018), and to perform identification based on individually distinct patterns (Wäldchen & Mäder, 2018; Weinstein, 2018). The work presented here investigates the use of deep learning methodologies to support semi-autonomous methods for sorting and identifying snow leopard individuals within an accessible format.

The Whiskerbook.org online platform (Whiskerbook 2021) provides a Web-based data management framework and a computer vision pipeline (Parham et al. 2018) for detection and individual identification of multiple species of large cats, including snow leopards. However, existing computer vision techniques on Whiskerbook.org, such as HotSpotter (Crall et al. 2013), have been recommended for use (Miguel et al. 2019) on snow leopards but not formally evaluated for accuracy, leaving questions about their overall accuracy and reliability for this species. Additionally, recent developments in machine learning have suggested that a new class of deep learning-based algorithms may improve automated matching capability (Moskvyak et al., 2019).

Researchers seek to address conservation and management for this species and conduct analyses over biologically relevant scales, meaningful to goals across the snow leopard range. The project needs to reconcile previously classified and curated snow leopard photo-ID catalogs with individuals from newly collected datasets. To do so, we need to automate a pipeline that is both more efficient in terms of expert time and reduces misidentification errors.

This work is novel as the first attempt at testing and thoroughly evaluating two computer vision algorithms to understand their performance at matching individual snow leopard sightings. The first is the Hotspotter algorithm (Crall et al. 2013), a SIFT-based comparison of significant visual texture areas, previously deployed on Whiskerbook.org for species like jaguar (*Panthera onca*). The second is called Pose Invariant Embeddings (PIE v2; Moskvyak et al. 2019), a convolutional neural network (CNN), in this case, InceptionV3 optimized with a triplet loss network, which was first tested on manta ray bellies (*Mobula* spp.) and humpback whale (*Megaptera novaeangliae*) flukes (Blount, 2018).

Understanding algorithm performance can inform the usability of the Whiskerbook.org platform and aid researchers in not only rapidly matching individuals across photo ID catalogs (a significant potential time and cost-saving on the road to more extensive and more comprehensive catalogs and modeling efforts) but also at improving the effectiveness of markre-capture models that ultimately inform snow leopard conservation.

## Materials and methods

### Camera Trap Imagery

We conducted our experiments and evaluations with curated photos of well-known individuals photographed between 2012 and 2019 from the Wildlife Conservation Society (WCS) program in Afghanistan, as recorded in the Whiskerbook.org platform (Blount, 2021). Additional data for captive snow leopard data were contributed from seven European zoos (Helsinki and Ätheri Zoos in Finland, Kolmården Zoo, Nordens Ark and Orsa Bear Park in Sweden, and Köln and Wuppertal Zoos in Germany) and two zoos in the United States (WCS managed Bronx and Central Park zoos in New York City). Our project had access to 22,120 annotations (i.e., machine learning-detected bounding boxes around snow leopards in photos) (Fig 1) from 359 snow leopard encounters within hourly intervals. However, these numbers are inflated by their data capture technique: camera traps, which generate a high volume of imagery in a brief timeframe and at a single location. For example, limiting the data to only individuals sighted three or more times (three annotations of a side is a minimum requirement for PIE model training) reduces the number of annotations to 12,311 and the number of distinct encounters to 116. Further data filtering used in machine learning training and analysis (e.g., to prevent overrepresentation of highly sighted individuals) reduced these numbers even further. While we believe this to be the largest data set yet assembled for analysis of computer vision on snow leopard individual ID, although the available data is still relatively small.

**Fig 1.**
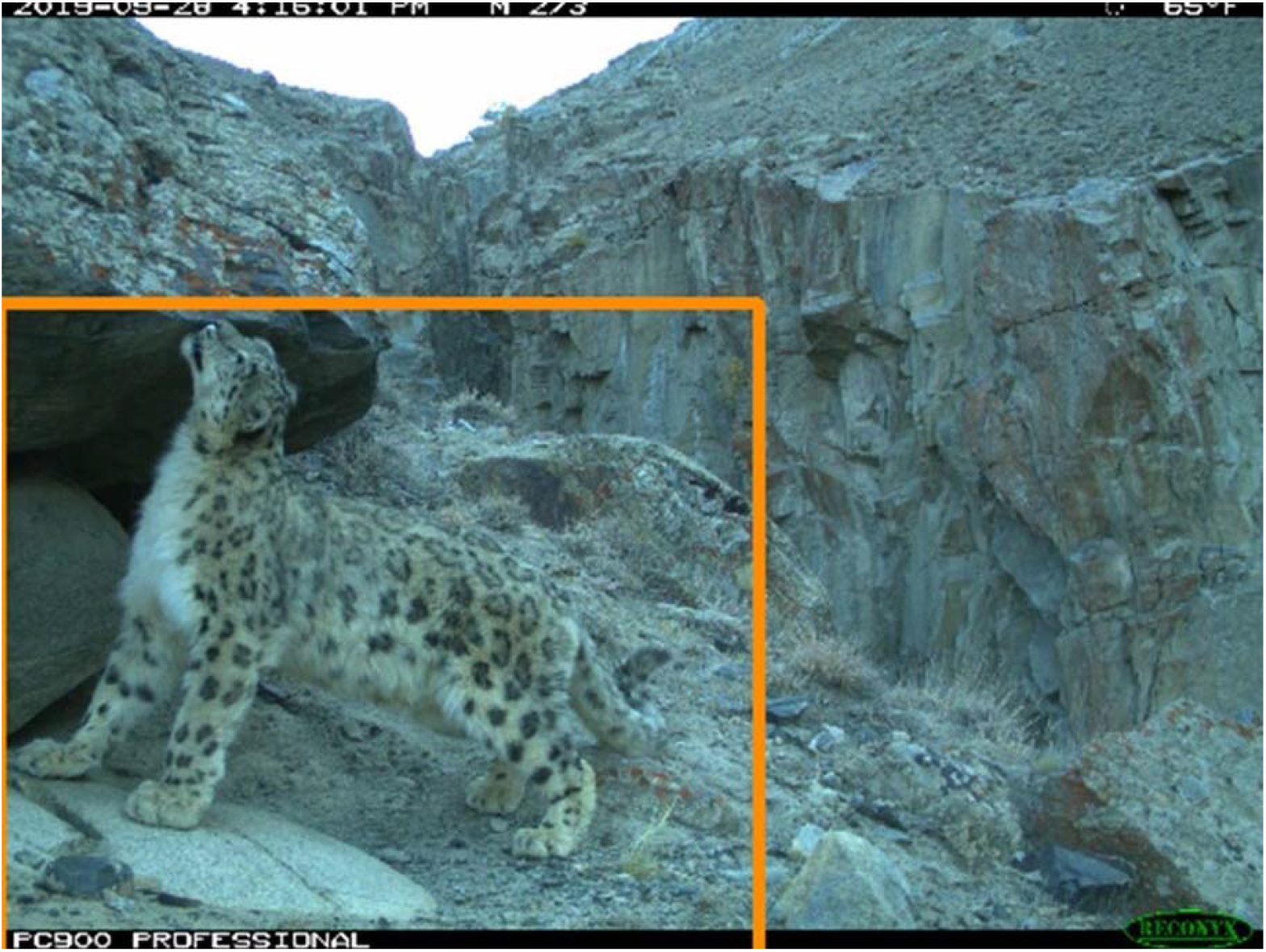
An annotated snow leopard. Annotations were generated by a machine learning-based computer vision model and associated with identifying known individuals by human confirmation. Annotations served as the fundamental data learned from ML and compared by each algorithm. Photo courtesy of WCS Afghanistan.

### Performance Metric

We evaluate the performance of each algorithm individually by computing the top-k accuracy on a test set where k = 1, 5, 10 and represents the rank of the correct match (i.e., an annotation of the same individual represented by a query annotation) in a list of proposed matches. A top-1 rank, therefore, is the correct result returned by the algorithm as the most likely match for a candidate annotation. A top-5 rank is the correct result ranked fifth most likely as returned in the candidate list and so forth.

#### Min-3/Max-10

For training machine learning algorithms like PIE, we often utilize a Min-3/Max-10 data subset. This data subset represents the individuals with at least three photos of the same viewpoint (either left or right), also limited to a maximum of ten photos per individual/viewpoint. The training phase requires a minimum of three photos (two photos for ML to learn from) and the test phase (at least one for ML to test it against). A maximum of ten photographs is allowed for data set balance and prevent highly sighted individuals from causing the ML system to optimize on highly sighted individuals yet perform poorly on infrequently sighted individuals. In our experience, a max-10 limit will suppress the Top-k performance ranking but create an ML model that performs better in real-world matching across various individuals. After applying these filters to the curated data on Whiskerbook.org, this resulted in 829 images of 217 individual snow leopards.

### Feature detection

Before the classification algorithms can act, the snow leopard needs to be detected in the images. A machine learning detector, a customized PyTorch implementation of YOLO v2 (Redmon et al. 2016), created the snow leopard annotations as the first step in the Wildbook Image Analysis (WBIA) pipeline (Parham et al. 2018). The task of the detector localizes animals in images, focused mainly on accurate bounding boxes over the ground-truth detections (made *a priori* by humans for a test set) while minimizing false positives and false negatives. We trained a model to predict the snow_leopard class (a species-labeled bounding box) using a training dataset of 2,000 images and 2,078 annotated bounding boxes (2 empty images, 34 images with two boxes, and 23 images with two boxes).

### Network architecture, data pre-processing, training, and evaluation

With a machine learning detector trained and configured to extract snow leopard annotations, we then used those annotations and related metadata (in particular the known identifications based on coat patterning) in the WBIA pipeline (Parham et al. 2018) to first custom train the Pose Invariant Embeddings (PIE) algorithm (Moskvyak et al. 2019).

We used a Min-3/Max-10 data constraint for PIE ML training and divided the training and test data (Table 1).

**Table 1.** Initial data division for machine learning training with PIE.

| Set | Individuals | Annotations |
| --- | --- | --- |
| train | 91 | 745 |
| test | 35 | 84 |
| total | 126 | 829 |

Models were trained and tested by tuning the number of required epochs and further assessing for any indication of overfitting based on model outputs and error results. Overfitting may occur if the model becomes too detail-oriented, fitting the data precisely, thus modeling extraneous noise in the training data instead of the general features of interest.

After performing the first round of PIE algorithm modeling, we observed PIE converging extremely quickly while training on this data. We believe this is partly due to territoriality (Only one location may have snow leopards photographs) and that much of the training data was from camera traps (compared to captive zoo data). We theorized that these two factors resulted in “background matching” as an effective strategy for the PIE model during training, essentially recognizing each snow leopard by recognizing the scenery where it most often appeared.

Researchers investigated background matching as a potential cause of overfitting by modeling two datasets, one using a subset of individuals (n=14) that occurred at multiple locations and a subset of individuals, including those that also had multiple sightings at the same location (n=29).

### Background Matching

We used two methods to minimize potential overfitting due to background matching in the PIE model developed during training. The first method is appropriately named “background subtraction” and removes the background algorithmically. Wild Me had already trained a background subtraction model for snow leopards as part of detector training in the WBIA pipeline (Parham et al. 2018), so we modified the PIE training pipeline so that each training image was pre-processed with background subtraction (Fig 2).

**Fig 2.**
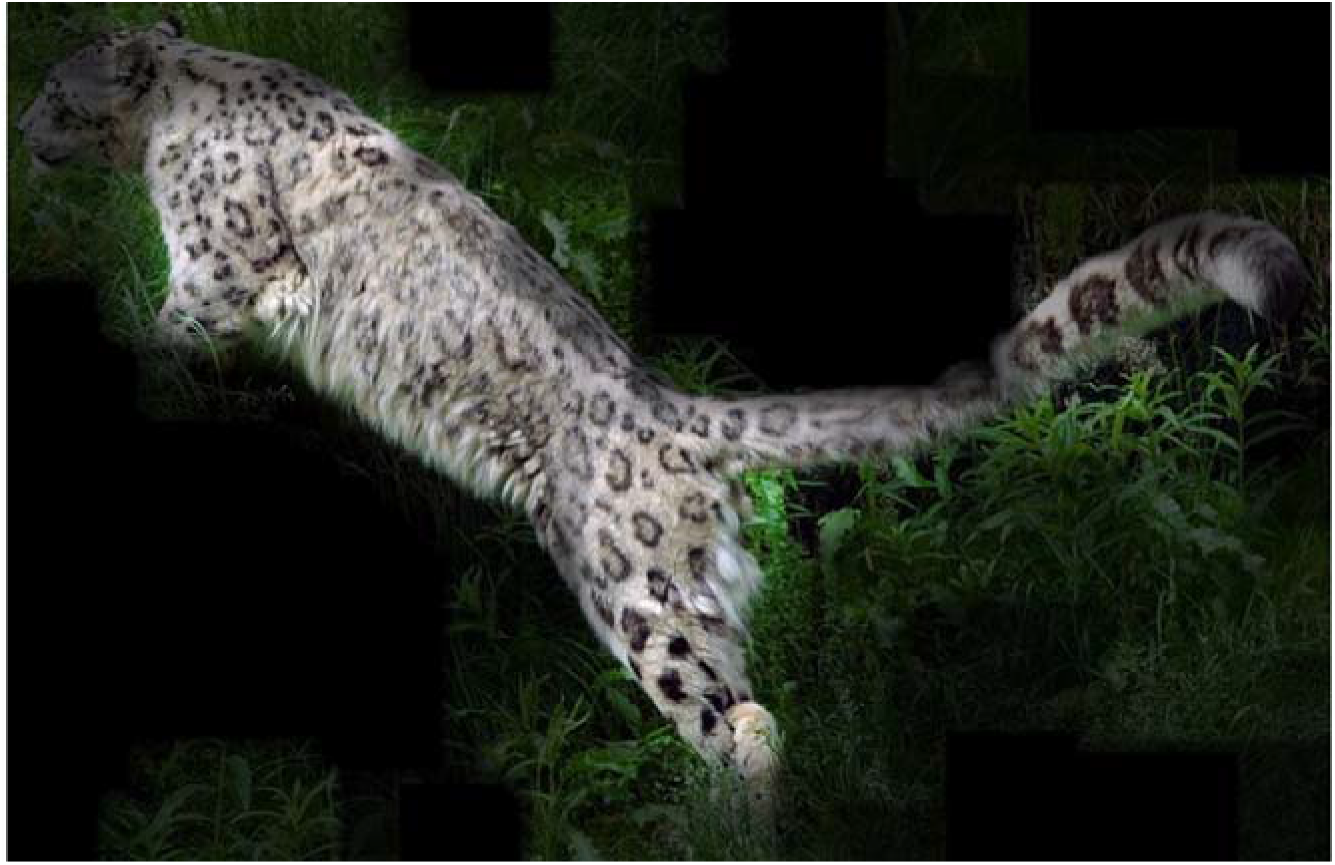
A background-subtracted snow leopard photo used to train the final PIE model. Credit: Wild Me

The second method involved mirroring left-side photos so that each image appears to have a right-side viewpoint of the animal. This mirroring is a configuration parameter while training PIE that can be turned on or off. It has previously been used on complicated matching problems to get as much image-level consistency as possible: PIE is generally able to match diverse viewpoints, but standardizing them may also increase accuracy.

## Results

### Detection Algorithm

The Precision-Recall performance curves of the trained snow leopard detection model were computed on a held-out 20% test set (413 annotations) to assess the accuracy and report comprehensive detection success (Fig. 3). The various colors of the curves show different thresholds of non-maximum suppression (NMS) applied to the network’s final bounding box predictions. A common way to summarize the localization accuracy is with Average Precision (AP) as determined by the area-under-the-curve. For example, the best performing configuration with an NMS of 30% achieves an AP of 94.45%. The corresponding colored points on each curve signify the closest point along the line to the top-right corner of the precision-recall coordinate system, signifying a perfect detector. Furthermore, the yellow diamond specifies the highest precision for all configurations, given a desired fixed recall of 80%.

**Fig 3.**
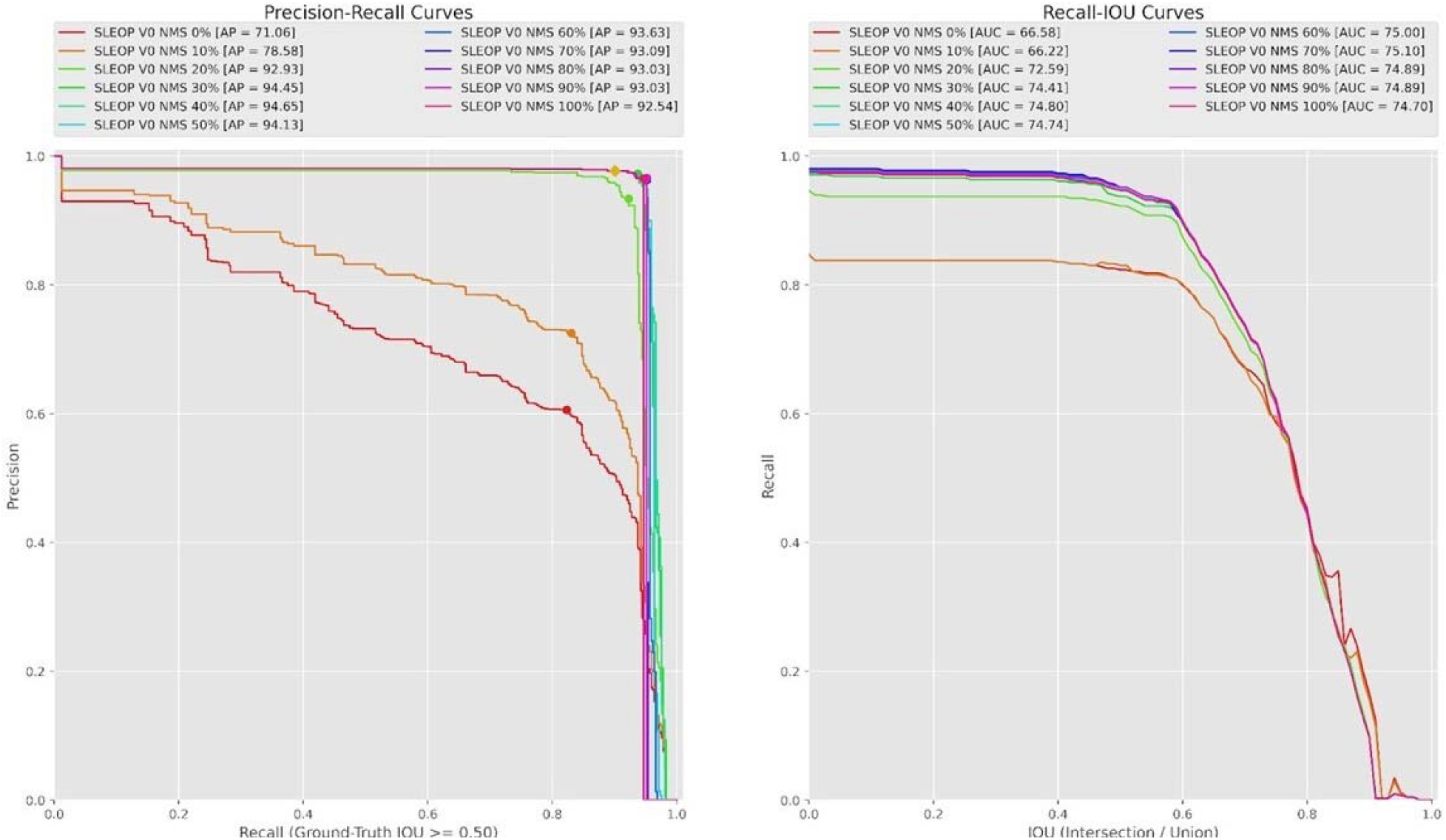
The detector Precision-Recall curves for snow leopards.

Delving deeper on the Precision-Recall curves, the maximum recall values (x-axis intercept) represent the absolute maximum percentage of annotations that the detector configuration can “recover” or “recall” from the ground-truth detections. Therefore, a recall of 90% indicates that a given detection configuration found 90% of the ground-truth annotations. The recall is a fundamental measurement for false negatives and implies a miss rate of 10% per sighting. The precision value indicates the percentage of correct detections (thereby measuring the number of false positives) and how many additional incorrect detections. A true-positive in our detection scenario is defined by the amount of intersection-over-union (IoU) percentage between a prediction and a matched ground-truth bounding box. For all plots in this section, we fix the acceptable IoU threshold to be 50% or greater. Non-maximum suppression (NMS) is a common technique for filtering duplicate detections by eliminating highly overlapping and lower-scoring predictions. A high NMS value will remove many bounding boxes from the output based on their percentage of overlap area (leading to an increase in precision but a decrease in recall). True negatives are undefined, which is why a receiver operating characteristic (ROC) curve is not provided in this report.

The confusion matrices give the accuracy for the best-colored point (left) and the yellow diamond (right) (Fig 4). It is worth mentioning that the 80% recall is arbitrary and can be adjusted based on the performance targets of the final project.

**Fig 4.**
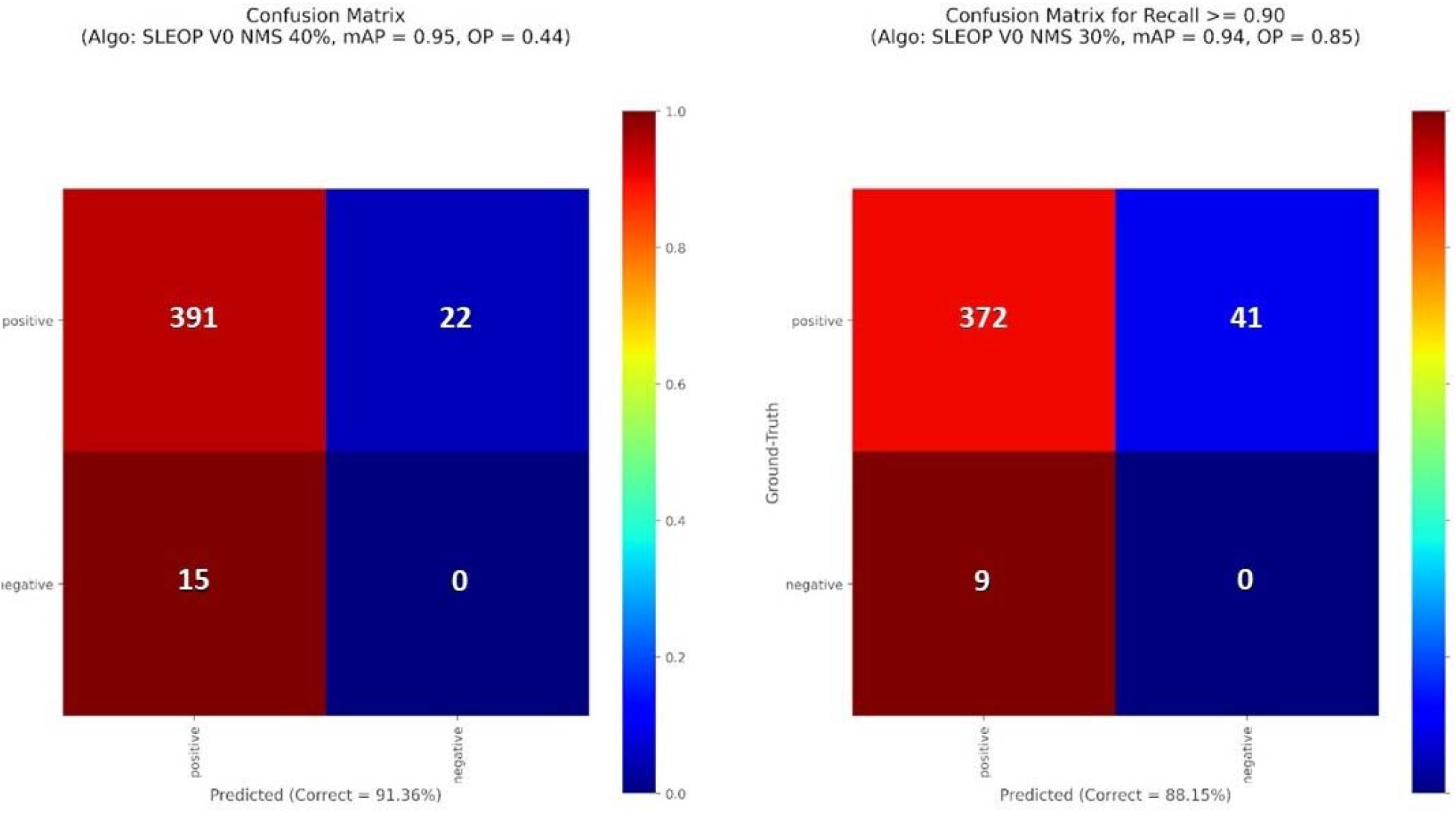
Confusion matrices for the best-colored point (left) and the yellow diamond (right) from the Precision-Recall performance curves (Fig. 2). False negatives occur when not detecting a snow leopard when one is present in the image, and false positives are spurious detections when no snow leopard is present in the image, such that a bounding box is generated where there is no snow leopard within it.

We can see that the best performing and our chosen configuration (highest AP at nearly 95%) has an NMS threshold of 40% and a score threshold of 44% (Fig 3 and 4). With this configuration, the overall detector makes 32 errors out of 413 overall annotations, with 22 of those incorrect detections being false negatives (not detecting a leopard when one is present). For 22 false negatives, there are 15 false positives (spurious detections of snow leopards that were not in the image, a bounding box is generated where there is no animal) (Fig 3). If we relax the miss-rate requirement to 10%, we make fewer false detections (a total of 9 down from 15), but we end up missing 41 animals (false negatives), and the overall accuracy drops by over 3% (Fig. 4, right plot).

Manual observers can remediate false positives and negative errors by cleaning the detection algorithm results. The cleaning can simultaneously deal with errors due to two detections being formed on the same individual, separate annotations creating new encounters for a second or third snow leopard or missing annotations. From user experience accounts, field camera trap data often contain many images of the same individual within an hourly interval, where the impacts of false negatives are likely not as significant on a larger dataset. For example, the detection algorithm may not classify several photos within an encounter of 30 photos, and the annotated sample for that individual is still substantial. Experienced observers report that false negatives seem to arise from low-quality captures, such as those inordinately far away from the camera trap or bad quality captures that are blurry or less recognizable.

### Investigation of Overfitting

After running an initial set of PIE models, the neural network reached its most accurate state rapidly, after seeing each image only 10-30 times (in machine learning terms, after 10-30 “training epochs”), whereas 70-250 epochs would be a more usual period for this convergence. Subsequent training only decreased the algorithm’s performance on held-out test data, meaning its long-term behavior was more akin to memorizing its training set than learning a generalized matching strategy (this is the machine learning definition of “overfitting”).

We investigated overfitting by measuring the algorithms’ accuracy matching 14 individuals which were seen at multiple locations (PIE results: Rank 1-68%, Rank-5 86%), and comparing with the initial model where there were also 29 individuals seen at the exact location with the same background (PIE results: Rank 1-77%, Rank-5 93%, and Rank-20 96%) (Fig 5). The algorithms’ top-1 and top-5 accuracies for snow leopard individuals detected at different locations are lower than the first modeling attempt, which we can determine were due to the background assisting in the classification of the image. These results clearly show the significance of background matching in increasing classification accuracy, despite the filtering criteria having used only half as many individuals across multiple locations. Since there were fewer individuals, we expected the individuals to classify with higher accuracy if the algorithm successfully identified the individual snow leopard in the image.

**Fig 5.**
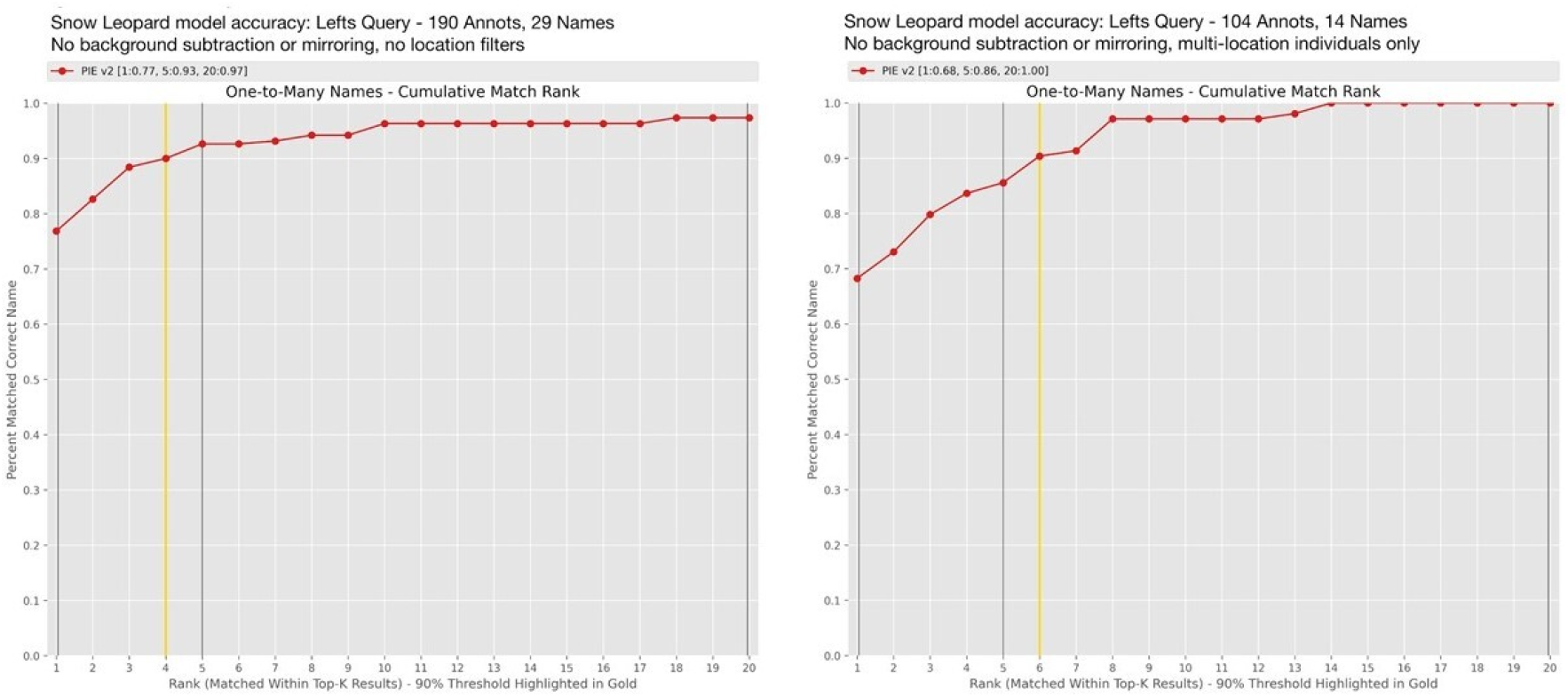
Accuracy on a PIE model without background subtraction or L/R mirroring. Left is without location filtering; right is only multi-location individuals that moved between trapping stations.

### Background Subtraction

The deployed model was then trained using imagery subjected to background subtraction and L/R mirroring. We found that turning on the LR parameter significantly changed the training behavior, causing slower convergence with fewer signs of over-fitting. We have speculated that this mirroring could double the number of background textures available for background matching. It is also possible that the slower convergence was due to the randomness in the initial configurations of the neural network during each training run.

We believe the PIE model with the mirrored and background-subtracted model has theoretical advantages and has shown higher accuracy, so we have chosen it for deployment in the Whiskerbook platform.

#### Min-2 Accuracies

The results shared so far are on datasets with at least three photos per individual + viewpoint (e.g., individuals with at least three left side photos), here referred to as “min-3 accuracy”. We also computed accuracies on the more conservative filter of “min-2”, which is the minimum required for a human reviewer to perform ID. These results include the scenario where the algorithm matches a new animal for the first time against only one existing catalog photo. The results sought to classify 40 individuals with PIE BGS-LR (Rank-1 72%, Rank-5 89%, Rank-20 94%), PIE without background subtraction (Rank-1 64%, Rank-5 86%, Rank-20 95%), as compared to Hotspotter (Rank-1 69%, Rank-5 82%, Rank-20 84%). The results showed that the PIE model with background subtraction performed the best on the min-2 side-by-side matching (Fig. 6). Overall, the BGS-LR model performs slightly better when compared to other models.

**Fig 6.**
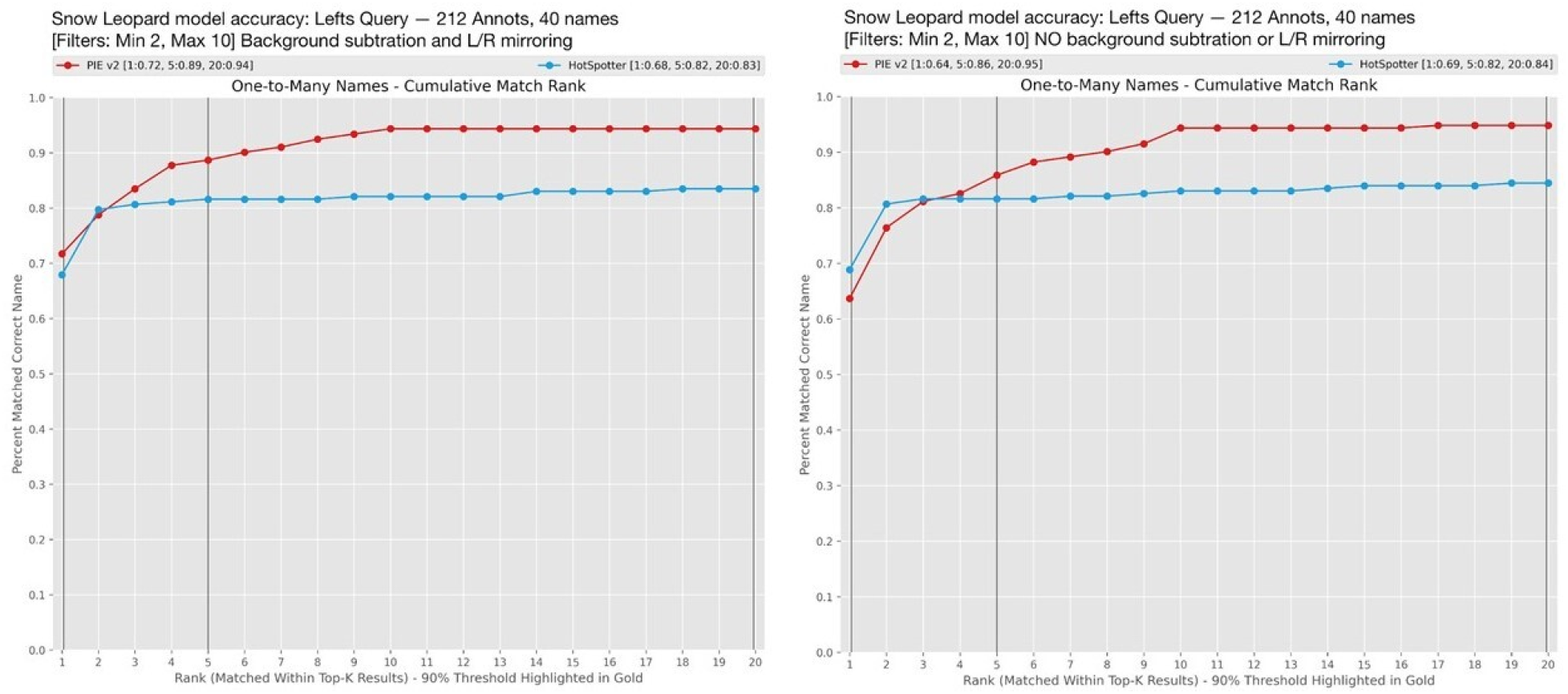
Comparison of our two PIE models on data with a minimum of two left-side photos per individual. The model on the left used background subtraction and L/R mirroring; the model on the right used neither technique. HotSpotter (HS) is shown in both cases; differences in HS accuracies are due to random noise as HotSpotter match scores are not strictly deterministic.

### PIE and Hotspotter Model Accuracy

Since Whiskerbook.org contains a multi-species, multi-feature, and multi-algorithm technical foundation (Blount et al., 2021), more than one algorithm can be run in parallel when identifying the individual animal in a photo. Therefore, we plotted the best PIE model alongside the older HotSpotter algorithm’s performance (Fig. 7), as well as the accuracy of both algorithms combined (i.e., top-1 of PIE + HotSpotter means the percentage of cases where at least one of the algorithms found the correct match at the top rank). Combining the two algorithms significantly improves overall match accuracy:

- Top-1: 85%
- Top-5: 95%
- Top-20: 99%

**Fig 7.**
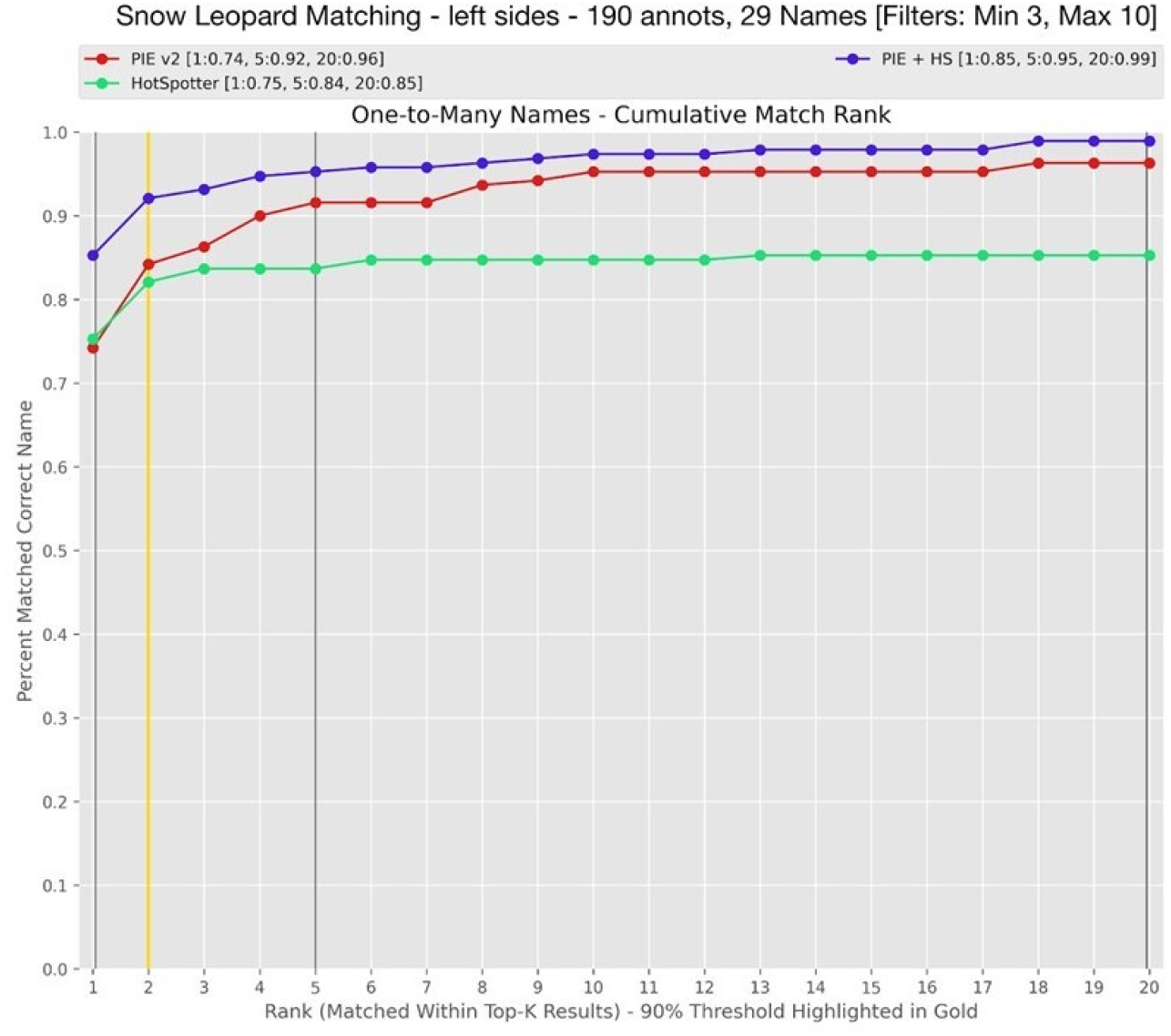
Accuracy plot of the deployed matching algorithm. Left-side annotations of individuals contain at least three left-side photos. Right-side performance is roughly equivalent (this model mirrors left-side photos, so every animal it sees is from a right-side perspective, but queries are pre-filtered by viewpoint not to compare lefts and rights).

The accuracy plots from all modeling attempts were summarized and compared across trained PIE model variations (Table 2). Two things determine the accuracy of an ID algorithm: the algorithm and the data. We have compared two PIE models, one trained with background subtraction and L/R mirroring (the final, deployed model, dubbed “PIE-BGS-LR” here), and one without either option (“PIE-vanilla”), as well as HotSpotter. We have considered three subsets of the same data, all identified snow leopard photos on Whiskerbook.org. The subsets are either min-3 or min-2, where the numeral indicates the minimum number of left-side sightings of the same individual required for those photos to be included in the subset (right-side matching behavior is comparable but not shown here). Additionally, datasets labeled “multiloc” indicate that the researchers filtered the data to only include animals seen at multiple locations while satisfying the min-X requirement.

**Table 2.** Accuracy summary of algorithms on various data filters. BGS stands for “Background Subtraction”. “LR” stands for left-right mirroring. PIE-Vanilla indicates unmodified annotation used in a PIE model. *: NA accuracies are shown where the result would be a trivial 100% because fewer than 20 individuals meet the criteria.

| Algorithm | Dataset | Number of Annots. | Number of Individuals | top-1 ACC | top-5 ACC | top-20 ACC |
| --- | --- | --- | --- | --- | --- | --- |
| PIE-BGS-LR | min-2 | 212 | 40 | 72% | 89% | 94% |
| PIE-vanilla | min-2 | 212 | 40 | 64% | 86% | 95% |
| HotSpotter | min-2 | 212 | 40 | 69% | 82% | 84% |
| PIE-BGS-LR | min-3 | 190 | 29 | 74% | 92% | 96% |
| PIE-vanilla | min-3 | 190 | 29 | 77% | 93% | 96% |
| HotSpotter | min-3 | 190 | 29 | 75% | 84% | 87% |
| PIE-BGS-LR + HotSpotter | min-3 | 190 | 29 | 85% | 95% | 99% |
| PIE-BGS-LR | min-3, multiloc. | 104 | 14 | 67% | 90% | NA* |
| PIE-vanilla | min-3, multiloc. | 104 | 14 | 68% | 86% | NA* |
| HotSpotter | min-3, multiloc. | 104 | 14 | 72% | 79% | 85% |

## Discussion

Our results provide a first look at the matching accuracy of two independent, production-ready pattern-matching algorithms for snow leopards. The results demonstrate that their use in combination can further improve researchers’ ability to identify matching snow leopards across large catalogs rapidly. Combining PIE and Hotspotter for classification yielded an accuracy of Rank 1-85% correctly identified individuals on our dataset. As individual algorithms, PIE has shown to have higher accuracy in matching these data than HotSpotter, and it is currently the best-performing algorithm we are aware of for identifying snow leopard individuals in camera trap photos. However, HotSpotter’s results are only slightly less performant, and since they are independently obtained, they add value in concert with PIE within the program.

Previous attempts at using deep learning for snow leopards advanced the methods to automatically detect snow leopards in photos (A. Miguel et al., 2016) and optimize classification using background erasing to assist Hotspotter algorithms to focalize on regions of interest (Beery, 2016; A. C. Miguel et al., 2019). These prior advancements improved our understanding of snow leopard individual identification capabilities and seeded the formal evaluation and proliferation of tools based on these successes.

One shortcoming identified and addressed within the study showed a limited background matching for the PIE model (Fig 8 helps visualize the impact). Rapid convergence indicated that the “matching problem” was abnormally easy on this data, which is generally a result of insufficient volume of training data, lack of diversity, or both. Background subtraction and left-right mirroring were included in the PIE algorithm model testing to resolve these issues.

The issue of background is significant for the snow leopard, as a territorial species, where field observations at one camera trapping site sometimes reveal many animals. Observers have reported two territorial males, two territorial females, and several younger cats at a single camera trapping site. Matching several individuals would be most effective when the algorithm is making intelligent ID predictions based on the natural patterning of the animal. However, if the model filters the imagery and makes intelligent decisions on the landscape features, it is not always a bad thing, where there may be minor effects that filter and rank the list of candidate individuals based on some background and location information.

These location-based limitations arise from compiling results based on a relatively small dataset from collated zoo data and field camera trap observations from one location in Afghanistan. Rapid convergence during PIE training suggests that more extensive and more diverse data may produce a more robust model. The snow leopard-PIE training pipeline developed here can be reused to train future models comparatively easily after more users begin to use the system and additional data can be then utilized. The existing model may also help bootstrap that data-curation process. There is a significant opportunity for regional and global-scale research collaborations with snow leopard research institutions to curate and individually identify data that would build on these existing models towards more sophisticated refinement and better performance.

We expect the PIE model deployed to Whiskerbook.org to save time during snow leopard matching, especially considering related Whiskerbook.org features. Features currently available include settings that allow for a location-filtering option (where users can limit a match query to animals sighted in a certain area), one-to-one image matching or one-to-many matching (for previously classified individuals), side-by-side comparison with HotSpotter results, and the overall high accuracy of PIE on our existing datasets documented here. The algorithm tools within Whiskerbook are also complemented by a “visual matcher” interface for manual classification by an observer, allowing for more easy side-by-side comparison of photos against each camera trap station’s photos at different dates.

Future research may seek to assess further the manual observer’s ability to classify images utilizing the features (Visual Matcher, Hotspotter, and PIE) within Whiskerbook using captive snow leopard individuals from zoo collected data. A study of this design could advance the results that demonstrated high misclassification rates from manual human labor on a set of captive snow leopards with known identities (Johansson et al., 2020). Johansson et al. (2020) showed that manual observers correctly classified 87.5% of all capture occasions, where misclassification errors would compound to inflate population abundance estimates 33% above actual population size. The algorithms demonstrated in this paper perform nearly at the level we would expect of a human observer, and the algorithms can serve as artificially intelligent observers and speed up the classification pipeline. Future research may seek to hypothesize and demonstrate manual observers’ increased capabilities using artificial intelligence and features afforded in Whiskerbook to bridge the accuracy gap towards more precise estimates.

## Data Availability

Research-related requests for annotations and data used for ML training in this paper can be requested in COCO format (Lin et al. 2020) via the corresponding author and must be expressly and independently permitted by author Eve Bohnett or through an established collaboration on Whiskerbook.org. Data can also be reviewed and shared via a collaboration request to user Eve Bohnett inside the Whiskerbook.org system.

## Code Availability

All software used in this analysis is available in the Wild Me open source repository at: https://github.com/wildmeorg

The base application for algorithm analysis as defined in Parham et al. 2018 is: https://github.com/WildMeOrg/wildbook-ia

Specific algorithm plugins for the three algorithms evaluated here can be found at: https://github.com/WildMeOrg/wbia-plugin-pie-v2

## Acknowledgments

This study was supported by the Global Environment Fund (GEF) and the United Nations Development Programme (UNDP)-grant AA/Pj/PIMS: 00105859/00106885/5844; project ‘Conservation of snow leopards and their critical ecosystems in Afghanistan’ and the European Union project “Improving participatory management and efficiency of rangeland and watershed focusing on Wakhan, Yakawlang, Kahmard and Sayghan Districts (Contract ACA/2018/399-742)” executed by the Wildlife Conservation Society (WCS) in Afghanistan. The paper’s contents are the sole responsibility of the authors and do not necessarily reflect the views of the European Union. We thank Patrick Thomas, WCS, and Craig Piper, WCS, for sharing photographs of captive snow leopards from Bronx and Central Park zoos, NY, USA. Special thanks to Tom Hoctor at the Center of Landscape Conservation Planning and Dave Hulse, the Florida Institute for Built Environment Resilience at the University of Florida. Machine learning advancements used in this study were partially funded by a Gordon and Betty Moore Foundation grant. A Microsoft Sponsorship supported Azure-based development and model deployment in Whiskerbook.org. The National Science Foundation partially funded this research under the Dynamics of Coupled Natural and Human Systems program [BCS-1826839]. This research also received financial and research support from San Diego State University.

## Author contributions

DB co-wrote and performed the PIE algorithm training and multi-algorithm analyses. JH co-wrote, fundraised, and coordinated data annotation for the project. JP trained the machine learning detector used before PIE training. SP, OJ, and SO collected the data. EB curated the data set used for machine learning training and performance analysis and performed writing, copy editing, and revising tasks. BA and WK contributed to drafting and revisions. LA contributed significant intellectual content. All authors provided comments and final approval of the uploaded manuscript.

## References

1. Beery, S. (2016). Orientation Invariant Autonomous Recognition of Individual Snow Leopards. 9.

2. Beery, S., Morris, D., Yang, S., Simon, M., Norouzzadeh, A., & Joshi, N. (2019). Efficient Pipeline for Automating Species ID in new Camera Trap Projects. Biodiversity Information Science and Standards, 3, e37222. https://doi.org/10.3897/biss.3.37222

3. Blount, D., Minton, G., Khan, C., Levenson, J., Dulau, V., Gero, S., Parham, J., & Holmberg, J. (2018). Flukebook – Continuing growth and technical advancement for cetacean photo identification and data archiving, including automated fin, fluke, and body matching. 13.

4. Blount, D., Parham, J., & Holmberg, J. (2021). Whiskerbook.org. Wild Me. http://www.whiskerbook.org/

5. Borchers, D., & Fewster, R. (2016). Spatial Capture–Recapture Models. Statistical Science, 31(2), 219–232. https://doi.org/10.1214/16-STS557

6. Choo, Y. R., Kudavidanage, E. P., Amarasinghe, T. R., Nimalrathna, T., Chua, M. A. H., & Webb, E. L. (2020). Best practices for reporting individual identification using camera trap photographs. Global Ecology and Conservation, 24, e01294. https://doi.org/10.1016/j.gecco.2020.e01294

7. Crall, J. P., Stewart, C. V., Berger-Wolf, T. Y., Rubenstein, D. I., & Sundaresan, S. R. (2013). HotSpotter — Patterned species instance recognition. 2013 IEEE Workshop on Applications of Computer Vision (WACV), 230–237. https://doi.org/10.1109/WACV.2013.6475023

8. Davis, R. S., Stone, E. L., Gentle, L. K., Mgoola, W. O., Uzal, A., & Yarnell, R. W. (2021). Spatial partial identity model reveals low densities of leopard and spotted hyaena in a miombo woodland. Journal of Zoology, 313(1), 43–53. https://doi.org/10.1111/jzo.12838

9. Ellis, A. R. (2018). Accounting for Matching Uncertainty in Photographic Identification Studies of Wild Animals [University of Kentucky Libraries; PDF]. https://doi.org/10.13023/ETD.2018.026

10. Falzon, G., Lawson, C., Cheung, K.-W., Vernes, K., Ballard, G. A., Fleming, P. J. S., Glen, A. S., Milne, H., Mather-Zardain, A., & Meek, P. D. (2019). ClassifyMe: A Field-Scouting Software for the Identification of Wildlife in Camera Trap Images. Animals, 10(1), 58. https://doi.org/10.3390/ani10010058

11. Foster, R. J., & Harmsen, B. J. (2012). A critique of density estimation from camera-trap data: Density Estimation From Camera-Trap Data. The Journal of Wildlife Management, 76(2), 224–236. https://doi.org/10.1002/jwmg.275

12. Jackson, R. M., Roe, J. D., Wangchuk, R., & Hunter, D. O. (2006). Estimating Snow Leopard Population Abundance Using Photography and Capture–Recapture Techniques. Wildlife Society Bulletin, 34(3), 772–781. https://doi.org/10.2193/0091-7648(2006)34[772:ESLPAU]2.0.CO;2

13. Johansson, Ö., Samelius, G., Wikberg, E., Chapron, G., Mishra, C., & Low, M. (2020). Identification errors in camera-trap studies result in systematic population overestimation. Scientific Reports, 10(1), 6393. https://doi.org/10.1038/s41598-020-63367-z

14. Mallon, Zhaler, P., McCarthy, T., Jackson, R., & McCarthy, K. (2016). IUCN Red List of Threatened Species: Panthera uncia. IUCN Red List of Threatened Species.

15. Miguel, A. C., Bayrakçismith, R., Ferre, E., Bales-Heisterkamp, C., Beard, J., Dioso, M., Grob, D., Hartley, R., Nguyen, T., & Weller, N. (2019). Identifying individual snow leopards from camera trap images. In K. Mao & X. Jiang (Eds.), Tenth International Conference on Signal Processing Systems (p. 36). SPIE. https://doi.org/10.1117/12.2521856

16. Moskvyak, O., Maire, F., Armstrong, A. O., Dayoub, F., & Baktashmotlagh, M. (2019). Robust Re-identification of Manta Rays from Natural Markings by Learning Pose Invariant Embeddings. ArXiv:1902.10847[Cs]. http://arxiv.org/abs/1902.10847

17. Nguyen, H., Maclagan, S. J., Nguyen, T. D., Nguyen, T., Flemons, P., Andrews, K., Ritchie, E. G., & Phung, D. (2017). Animal Recognition and Identification with Deep Convolutional Neural Networks for Automated Wildlife Monitoring. 2017 IEEE International Conference on Data Science and Advanced Analytics (DSAA), 40–49. https://doi.org/10.1109/DSAA.2017.31

18. Norouzzadeh, M. S., Morris, D., Beery, S., Joshi, N., Jojic, N., & Clune, J. (2019). A deep active learning system for species identification and counting in camera trap images. ArXiv:1910.09716 [Cs, Eess, Stat]. http://arxiv.org/abs/1910.09716

19. Nyhus, P. J., McCarthy, T., & Mallon, D. P. (2016). Snow Leopards: Biodiversity of the World: Conservation from Genes to Landscapes. Elsevier Inc.

20. Parham, J., Stewart, C., Crall, J., Rubenstein, D., Holmberg, J., & Berger-Wolf, T. (2018). An Animal Detection Pipeline for Identification. 2018 IEEE Winter Conference on Applications of Computer Vision (WACV), 1075–1083. https://doi.org/10.1109/WACV.2018.00123

21. Redmon, J., & Farhadi, A. (2016). YOLO9000: Better, Faster, Stronger. ArXiv:1612.08242 [Cs], http://arxiv.org/abs/1612.08242

22. Royle, J. A., & Young, K. V. (2008). A HIERARCHICAL MODEL FOR SPATIAL CAPTURE–RECAPTURE DATA. Ecology, 89(8), 2281–2289. https://doi.org/10.1890/07-0601.1

23. Wäldchen, J., & Mäder, P. (2018). Machine learning for image based species identification. Methods in Ecology and Evolution, 9(11), 2216–2225. https://doi.org/10.1111/2041-210X.13075

24. Weinstein, B. G. (2018). A computer vision for animal ecology. Journal of Animal Ecology, 87(3), 533–545. https://doi.org/10.1111/1365-2656.12780

